# Reconfigurable microfluidic circuits for isolating and retrieving cells of interest

**DOI:** 10.1101/2021.12.23.473995

**Authors:** Cyril Deroy, James H. R. Wheeler, Agata N. Rumianek, Peter R. Cook, William M. Durham, Kevin Foster, Edmond J. Walsh

## Abstract

Microfluidic devices are widely used in many fields of biology, but a key limitation is that cells are typically surrounded by solid walls, making it hard to access those that exhibit a specific phenotype for further study. Here, we provide a general and flexible solution to this problem that exploits the remarkable properties of microfluidic circuits with fluid walls – transparent interfaces between culture media and an immiscible fluorocarbon that are easily pierced with pipets. We provide two proofs-of-concept in which specific cell sub-populations are isolated and recovered: i) murine macrophages chemotaxing towards complement component 5a, and ii) bacteria (*Pseudomonas aeruginosa*) in developing biofilms that migrate towards antibiotics. We build circuits in minutes on standard Petri dishes, add cells, pump in laminar streams so molecular diffusion creates attractant gradients, acquire time-lapse images, and isolate desired sub-populations in real-time by building fluid walls around migrating cells with an accuracy of tens of micrometres using 3D-printed adaptors that convert conventional microscopes into wall-building machines. Our method allows live cells of interest to be easily extracted from microfluidic devices for downstream analyses.

## Introduction

Microfluidic experiments are widely used to study the physical and chemical stimuli that influence individual cell behaviour and physiology, and have advanced understanding in both prokaryotic and eukaryotic biology [1]. While cells within microfluidic devices can be easily visualised by time-lapse microscopy, it remains difficult to retrieve responding cells from widely-used devices that are made with polydimethylsiloxane (PDMS), as they are confined behind solid walls. This limits our ability to relate the phenotypes observed within these systems with their underlying molecular processes. Although open microfluidics improves accessibility [2], it remains challenging to isolate and extract specific cells of interest from flowing micro-environments.

The recent development of fluid-walled microfluidics – where samples in the aqueous phase are confined behind a liquid interface made of the immiscible fluorocarbon, FC40 – allows fluid walls to be built by a three-axis traverse (a ‘printer’) [3]. This approach can be used to construct isolating fluid chambers around cells growing on standard Petri dishes [4], but it is difficult to do so around cells with specific phenotypes that are usually detected with a microscope. Furthermore, cells are often sensitive to changing flows [5]–[7], and so even moving a dish from microscope to printer can perturb both a cell’s position and physiology. Therefore, we devised a general method to isolate and extract cells from microfluidic circuits guided by traditional inverted microscopes and without interrupting flows.

As proofs of concept, we isolate and extract sub-populations from two very different cell types – murine macrophages and early-stage biofilms of *Pseudomonas aeruginosa* – as cells migrate up flow-generated gradients of two very different chemoattractants (i.e., macrophages towards complement component 5a (C5a), [8], [9], and the bacteria – counter-intuitively – towards the lethal antibiotic, ciprofloxacin [10]). We first show that fluid-walled circuits can be reconfigured using the printer to isolate a microscopically-targeted population of migrating macrophages. Secondly, we 3-D print adaptors that can be attached to a conventional inverted microscope with a motorized stage, transforming the microscope into a tool that is used to directly reconfigure the microfluidic device and isolate a sub-population of cells. This means that circuits can be reconfigured *in situ* on the microscope as flows and imaging continue. These methods are easily tuned to a user’s imaging system and requirements, giving them broad applicability to a wide range of experimental systems and questions in the biosciences.

## Results

### Workflow

The general workflow involves printing an ‘m’-shaped circuit with two arms that join to form a central conduit connected to a sink and seeding cells into the central conduit where they attach to the surface of the dish (**Fig. 1Ai**). Dishes are then transferred to a microscope where stable chemoattractant gradients are established over cells. Needles are inserted into each circuit arm, and medium infused through one and medium plus attractant through the other; a third needle inserted into the sink simultaneously withdraws fluid at a rate matching the two inputs. All three needles are connected to syringe pumps via tubing and held above the circuit by a 3-D printed holder attached to the dish. Both input liquids flow as laminar streams down the central arm, and diffusion creates a stable gradient of attractant across its width (**Fig. 1Aii**; **Fig. S1**; **Materials and Methods**). Once cells move up the gradient, new fluid walls are built around responding cells to isolate them from all others using either a printer or modified microscope (**Fig. 1Aiii**), and cells are then extracted prior to downstream analysis (**Fig. 1Aiv,v**).

**Figure 1.**
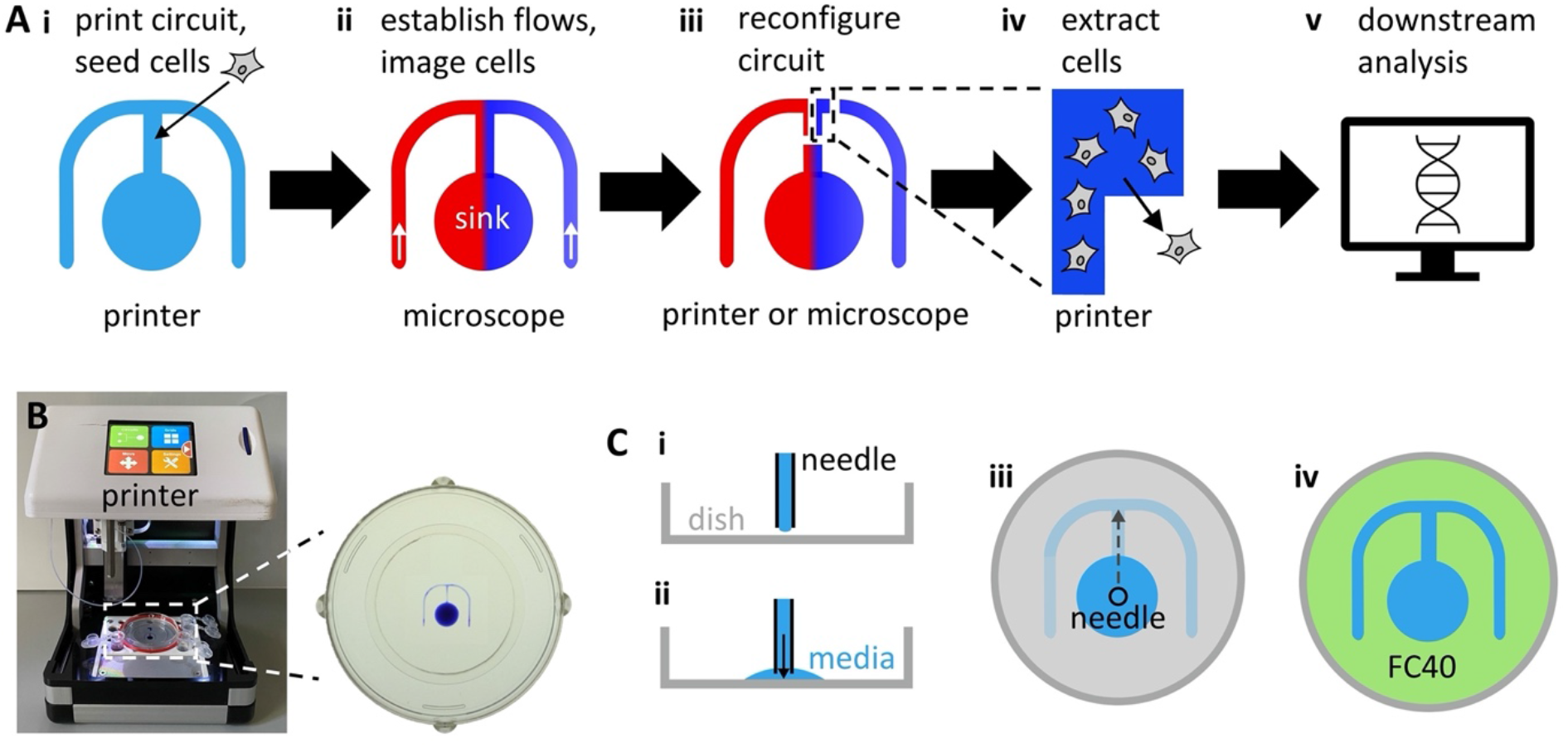
General workflow and printing circuits. **(A)** Workflow. **(i)** An ‘m’-shaped circuit is printed on a dish, and cells seeded into the central arm. **(ii)** The dish is transferred to a microscope, dispensing needles inserted into lateral arms of the ‘m’, and media -/+ chemoattractant (red/blue) pumped into the circuit to flow to the sink; diffusion between laminar streams creates a stable attractant gradient across the central conduit. The microscope is used to image chemotaxing cells and to identify rapidly-migrating ones. **(iii)** New isolating fluid walls are built around chemotaxing cells (on either the printer or microscope). **(iv)** The printer extracts isolated cells from the reconfigured chamber. **(v)** Cells are now analysed in any desired way (e.g., by RNA sequencing). **(B)** Printer with 6 cm dish. Close-up shows the dish with printed circuit (+ blue dye for visualization). **(C)** Printing the circuit. **(i)** A dispensing needle is held by the printer above a dish. **(ii)** Media pumped on to the dish is held in place by interfacial forces. **(iii)** Moving the needle as it infuses media prints the circuit. **(iv)** Overlaying FC40 (green) prevents evaporation.

#### Printing fluid-walled microfluidic circuits

The circuit is made using a custom printer that consists of a dispensing needle connected to a syringe pump mounted on a three-axis traverse (**Fig. 1B**; [3]); the needle infuses growth medium onto the surface of a standard Petri dish (**Fig. 1Ci-iii**, see **Materials and Methods**). Once printed, the circuit is overlaid with FC40 – a biocompatible and immiscible fluorocarbon – that prevents circuit evaporation (**Fig. 1Civ**). At the microscale, interfacial forces firmly pin medium to the substrate, and fluid walls – interfaces between the two immiscible phases – confine medium to the printed pattern. Fluid is easily added to and removed from such a circuit as fluid walls can be pierced at any point by pipets or dispensing needles; these walls spontaneously seal around inserted tubes, and then reseal without leaks when tubes are withdrawn. Fluid walls also morph above unchanging footprints to accommodate accompanying pressure differences (i.e., pinning of the contact line ensures that the circuit footprint remains unchanged as heights and contact angles change [3]). Soon after printing, pressure inside these circuits equilibrates causing the flow to cease. While we print these circuits by infusing medium through a needle, they can also be fabricated using a stylus or a micro-jet [11], [12].

#### Managing positional information

During the next and subsequent steps in the workflow (**Fig. 1A**), the dish is moved between printer and microscope, and the circuit’s positional information must be transferred between the two. This is achieved using 3-D printed adaptors that fix the orientation of the dish on the two instruments. Thus, a locating ring with a triangular protrusion attaches to the outside of the dish (**Fig. 2Ai**), and the ring and protrusion fit into appropriate indents in custom adaptors on the stages of the printer and microscope (**Fig. 2Aii,iii**).

**Figure 2.**
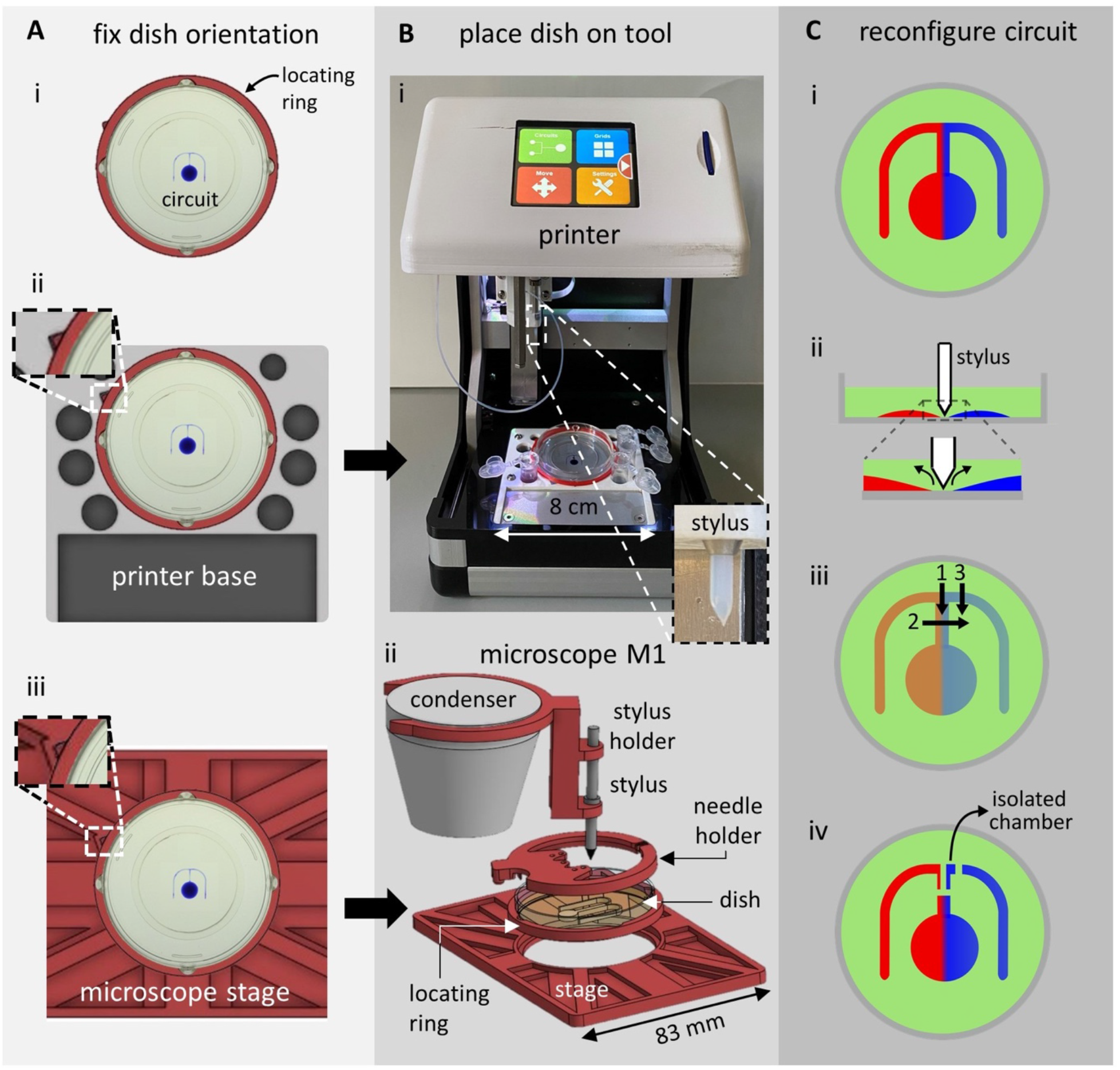
Reconfiguring microfluidic environments on a printer or microscope. **(A)** Fixing orientation of dish. **(i)** A 3-D printed locating ring (red) with a triangular protrusion (top left). The ring fits around a 6 cm dish containing a circuit (+ blue dye for visualization). **(ii)** The stage of the printer bears a complementary triangular indentation that orients the dish. **(iii)** The ring also fits into an adaptor for the microscope stage (red, top-down view). **(B)** Reconfiguration tools. **(i)** Printer with dish in red locating ring. Inset: close-up of PTFE stylus used to reconfigure circuit. **(ii)** Microscope condenser with exploded view of 3-D printed attachments (red; see **Supplementary STEP files** for CAD models). **(C)** Cartoons illustrating circuit reconfiguration. **(i)** Circuit before reconfiguration (red/blue: media -/+ chemoattractant). **(ii)** The stylus is lowered onto the surface of the dish; it forces media aside and brings FC40 down to wet the dish where it remains stuck to the bottom. **(iii)** The stylus builds new fluid walls along paths 1-3. **(iv)** This results in 3 new fluid walls that completely isolate part of the circuit from the rest.

#### Exposing cells to chemical gradients, imaging, and isolating migrating cells

After printing the circuits, dishes are transferred to a microscope, needles inserted, stable gradients of chemoattractant established, cells imaged as they migrate up the gradient, and the desired sub-population identified (**Fig. 1Aii**). This sub-population is isolated by building new fluid walls around it (**Fig. 1Aiii**) using a PTFE (polytetrafluoroethylene) stylus with a conical tip. This is achieved in two ways that use the same principle; the stylus is attached either to the three-way traverse on the printer (**Fig. 2Bi**, inset), or to the condenser on a standard inverted microscope with a motorized stage (**Fig. 2Bii**; the stylus is attached to the condenser using 3-D printed adaptors, **Supplementary STEP files**). In both cases, the stylus is lowered (either using the programmed traverse, or by lowering the microscope condenser manually) through FC40 and the medium in the circuit until the tip contacts the surface of the dish. As FC40 wets PTFE and polystyrene dishes better than medium [13], this brings the FC40 down onto the bottom of the dish. Next, the stylus is moved relative to the substrate, and – as it moves laterally – it displaces media from within the circuit. This displaced media is replaced by surrounding FC40, creating new fluid walls that can be constructed around the desired sub-population of migrating cells (**Fig 2C**).

Prior to reconfiguring circuits on the printer (**Fig. 2Bi**), flow through the circuit on the microscope is stopped, and input and output needles withdrawn. Owing to the adaptors (**Fig. 2A**), the dishes are then transferred back onto the printer in a fixed orientation. As macrophages adhere strongly to polystyrene and move at ∼1 µm min^-1^ at 37°C [14], the relative positions of responding and non-responding macrophages change little during transfer to the printer, or subsequently in the few minutes the printer takes to build new isolating walls at room temperature. However, a small proportion of bacterial cells often detach from surface-attached biofilms [15]–[17]. Under continuous flow, these planktonic cells are flushed through the circuit but any disturbance to flow could allow planktonic cells to swim into different parts of the circuit, where they could subsequently re-attach. To prevent this, we build the first fluid wall that separates chemotaxing bacteria directly on the microscope so that flows are not interrupted by having to move the circuit back onto the printer. In this case the stylus is fixed to the microscope condenser, and circuits are reconfigured as the motorised stage moves the dish relative to the stationary stylus. Then, new fluid walls are built as before in seconds with an accuracy largely determined by that of the microscope stage (typically in the micrometre range) and the thickness of the stylus tip (tens of micrometres). This setup therefore allows chemotaxing cells to be isolated from others without interrupting flows or imaging.

### Recovering macrophages chemotaxing towards C5a

Complement component 5a (C5a) is a protein that plays an important role in the innate immune response and acts as a chemoattractant that recruits macrophages to infection sites [18]. As a proof-of-principle of the first workflow outlined above where circuits are reconfigured on the printer, a population of murine bone-marrow-derived macrophages (BMDMs) are exposed to gradients of C5a (**Fig. 3Ai**; exact circuit dimensions shown in **Fig. S2A**), and the migrating sub-population isolated and extracted.

**Figure 3.**
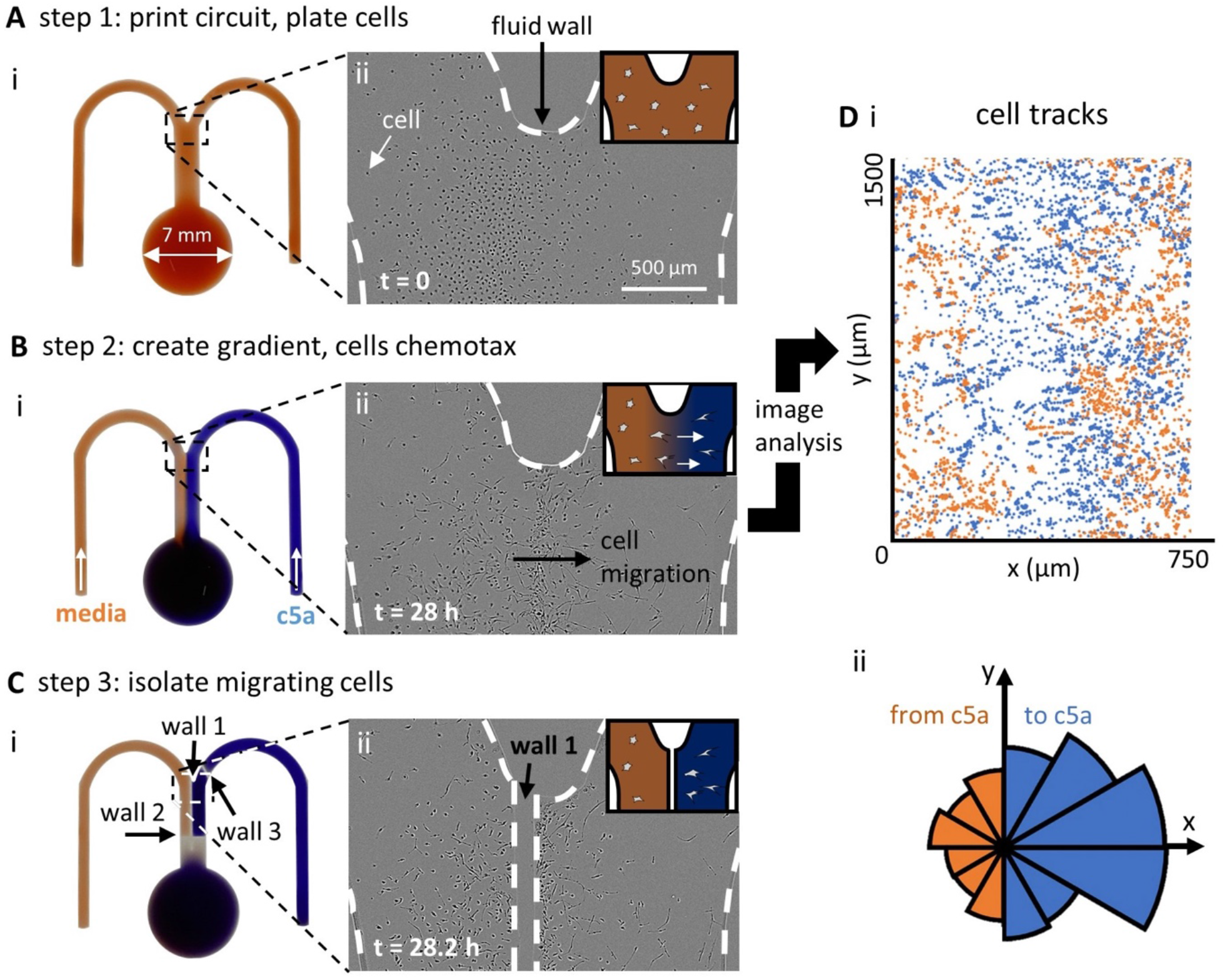
Steps in building fluid walls around chemotaxing macrophages (BMDMs). **(A)** Step 1 (after printing a circuit and plating BMDMs; *t* = 0 h). **(i)** Image of ‘m’-shaped circuit (with red dye to aid visualisation; dish invisible). **(ii)** Phase-contrast micrograph of area indicated in (A,i). Cells (dark dots) are confined by medium:FC40 walls (dashed lines). Inset: cartoon showing cells in media (red). **(B)** Step 2 (after establishing a chemoattractant gradient between laminar streams of media ± 10 nM C5a, which flow down the conduit at 1 µL min^-1^; *t* = 28 h). **(i)** Image of circuit. As before, red and blue dyes represent media and C5a. **(ii)** A micrograph of area in (B,i) taken from a movie in which many cells are seen to chemotax towards C5a on the right. Inset: cartoon showing cells chemotaxing towards C5a (blue). **(C)** Step 3 (after building three new fluid walls around chemotaxing cells on the printer; *t* = 28.2 h). **(i)** Image of circuit. New walls again faintly visible (positions also indicated by dashed white lines). Pumps are stopped before building new walls, and below wall 2 there has been time for diffusion to dissipate the dye gradient across the conduit. **(ii)** Micrograph of area in (C,i) showing “wall 1” built down the conduit separating chemotaxing from non-chemotaxing cells. **(D)** Image analysis of cells seen in movie. **(i)** Trajectories of individual cells in (bii) (blue – towards C5a, orange – away from C5a). **(ii)** Rose plot indicating binned probability-density distribution of directed cell movement (total number of trajectories *n* = 841, three experimental repeats). Movement is strongly biased towards C5a.

#### Establishing the gradient

After printing the circuit and seeding cells, input and output needles are inserted into appropriate positions in the circuit using a 3-D printed holder (**Fig. S1A,B**), and input (i.e., medium ± 10 nM C5a) and output flows established (**Fig. 3Bi**). Previous assays using dyes confirm that diffusion creates a stable concentration gradient across the width of the central conduit (left-hand inset in **Fig. 3Bi**; maximum gradient width is ∼53 µm – **Materials and Methods**).

#### Identifying the chemotaxing population

The dish is now placed in a commercially-available imaging system – an ‘IncuCyte’ ZOOM – that fits in a CO_2_ incubator; the microscope objective can be programmed to collect images as it moves under stationary dishes. Note, however, that our approach is easily extended (using appropriate 3-D printed adaptors) to most automated microscopes suitable for live-cell imaging of mammalian cells. In our case, cells in the IncuCyte are imaged for 28 h at 3 frames h^-1^. Real-time observation reveals that BMDMs chemotax towards C5a, as expected. **Figure 3Bii** illustrates one frame from **Supplementary Video 1**. Macrophages – initially randomly distributed throughout the conduit – accumulate approximately along the centre-line of the C5a gradient. To confirm directed movement, we analysed trajectories of 841 individual cells using custom particle-tracking software, (**Materials and Methods**). Where the C5a concentration gradient is steepest (close to the centre-line), there are many more cell trajectories moving towards C5a compared to the number moving away from the chemoattractant (shown in blue and orange, respectively, in **Fig. 3Di**), and we confirm this bias in a rose plot (**Fig. 3Dii**). Our results are consistent with a previous study that shows BMDMs robustly chemotax towards 10 nM C5a [8].

#### Isolating the chemotaxing sub-population

Migrating BMDMs are now isolated by constructing new fluid walls around them (**Fig 3C**). Flow through the circuit is halted and the dish returned to the printer where the stylus builds a new separating fluid wall (“wall 1” in **Fig. 3Ci**) down the centre-line of the conduit (**Fig. 3Cii**). Subsequently two additional walls are built across the width of the central conduit (“wall 2” and “3” in **Fig. 3Ci**) to create an entirely isolated fluid chamber (∼500 nL containing the sub-population of cells that migrated towards C5a, in addition to any non-responders initially present).

#### Extracting the chemotaxing sub-population

The printer now infuses ethylenediaminetetraacetic acid (EDTA) – a calcium chelator that detaches macrophages from surfaces [19] – into the chamber, and after 10 min, the aqueous phase containing now-suspended cells is withdrawn (along with a small amount of surrounding FC40) and manually plated into conventional tissue-culture dishes. Using this approach, ∼87% of the migrating cells that had been isolated in the fluid chamber are extracted and plated, of which ∼80% remain viable after 24 h (**Fig. S3**). This value is equivalent to those achieved with conventional cultures of these fragile primary cells [19]. These results confirm that a sub-population rich in migrating cells can be isolated from the general cell population with excellent viability. Where desired, this approach could easily be extended to carry out, for example, detailed downstream molecular analysis of chemotaxing and non-chemotaxing populations in order to connect phenotypical differences to variations in gene expression (**Fig. 1Av**).

### Recovering *P. aeruginosa* cells migrating towards antibiotics

*P. aeruginosa* cells are able to grow in a planktonic state but will also attach to surfaces and grow there as a biofilm. During the early stages of biofilm formation, surface-attached *P. aeruginosa* cells move by twitching motility using type-IV-pili to drag themselves across the substrate at speeds of ∼0.2-0.5 µm min^-1^ [20], [21]. Twitching *P. aeruginosa* cells undergo chemotaxis towards (and not away from) lethally-high concentrations of antibiotics [22]. In a companion paper, we characterise the biology of this response, which appears to have its basis in an aggressive response to toxins that come from competing bacteria [22]. Several factors made studying bacterial chemotaxis towards antibiotics challenging and thus this phenotype was an ideal test case for our approach [22]. First, a population of *P. aeruginosa* cells frequently detach during the earliest stages of biofilm development to become planktonic [15], where they use flagella to swim four orders of magnitude more quickly than surface-attached cells (at ∼2,400 µm min^-1^, [23]); consequently, they can traverse the microfluidic channels described above in seconds. However, the flow velocity in the laminar streams we use during experiments is ∼21,600 µm min^-1^, so ordinarily any cells becoming planktonic are rapidly flushed away. Second, the steep gradients required to detect chemotaxis in our circuits break down within minutes once flows cease. This would expose cells to rapid changes in antibiotic concentration that can harm them and cause them to detach from the surface. Consequently, the chemotaxing sub-population in a biofilm is best isolated without perturbing flow. Therefore, we designed a workflow to isolate chemotaxing cells under flow.

#### Imaging bacteria within antibiotic gradients

Here, we use a smaller ‘m’-shaped circuit with a narrower junction to limit the diffusion of antibiotic near the wall of the junction (**Fig. 4Ai**; **Fig. S2B**). As the chemical gradients in this device are a function of flow and molecular diffusion, a narrower junction ensures that antibiotic is not transported along the circuit walls, where the flow is minimal. *P. aeruginosa* cells are inoculated into the central conduit, where they attach to the dish and develop into an early-stage biofilm (**Fig. 4Aii**). Cells are seeded as a 1:1 co-culture of wild-type (WT) and mutant cells (*ΔpilG*) – the WT expresses the yellow fluorescent protein (YFP), and the mutant acts as an internal chemotaxis-null control as it lacks the *pilG* gene required to bias movement and undergo chemotaxis [21]. After mounting the dish on the microscope stage, media ± ciprofloxacin (an antibiotic often used clinically to treat *P. aeruginosa* infections) is infused into the circuit, and – as before – diffusion creates a stable gradient of attractant across the central conduit (maximum gradient width ∼70 µm; **Materials and Methods**).

**Figure 4.**
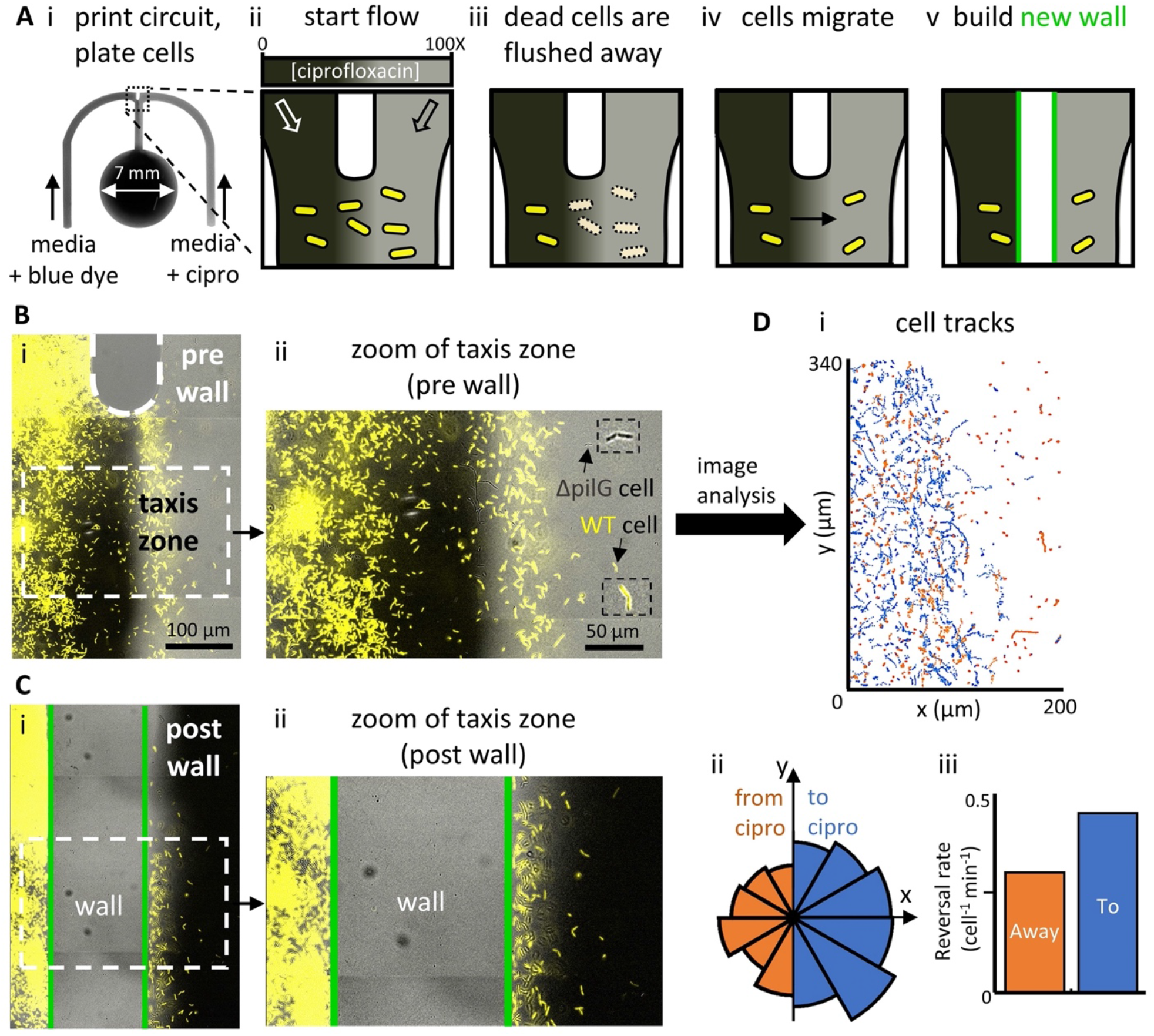
Building fluid walls on a microscope to isolate bacteria chemotaxing towards an antibiotic. **(A)** Overview. **(i)** Micrograph of ‘m’-shaped circuit. A 1:1 co-culture of YFP-labelled WT and unlabelled *ΔpilG* mutant (chemotaxis-null) *P. aeruginosa* are seeded in the conduit, and laminar streams of media + blue dye (dark grey) and media + ciprofloxacin (cipro, 10 μg mL^-1^ = 100X MIC; light grey) flow over them. **(ii-v)** Cartoons giving overview (grey scale reflects antibiotic concentration, only WT bacteria shown). After plating, WT bacteria grow in the conduit. Flow then establishes a concentration gradient by diffusion of antibiotic across the conduit; cells exposed to high concentrations on the right die and are washed away. Some bacteria then chemotax from the antibiotic-free region into the high concentration of ciprofloxacin. A new wall is then built between chemotaxing and non-responsive cells. **(B)** Micrographs of the top of the central conduit before building a new isolating wall. **(i)** Merge of fluorescence and phase-contrast images showing cells in a ciprofloxacin gradient (only fluorescent WT cells visible at this magnification). Most cells on the right have been killed and washed away, and some on the left are chemotaxing towards the antibiotic. **(ii)** Zoom of cells in taxis zone. The area is rich in yellow-fluorescing WT cells, with some phase-dark mutant cells (arrow and insets mark both types of cell). **(C)** Micrographs after building the wall. **(i)** Merge of fluorescence and phase-contrast images showing cells (only WT visible) and the new wall (cells within it are removed during building). The chamber on the right now contains ciprofloxacin at 3 times the MIC plus blue dye (dark grey, see **Materials and Methods**). **(ii)** Zoom of taxis zone and new wall. **(D)** Image analysis of cells in the taxis zone prior to building new wall. **(i)** Trajectories of individual cells (blue/orange tracks – to/from antibiotic. **(ii)** Rose plot indicating binned probability density distribution of directed movement (total number of trajectories n = 1470; 3 experimental repeats). Movement is biased to the right. **(ii)** Cells bias motility towards antibiotic by reversing direction more frequently when moving away from the antibiotic.

For these experiments, one input contains ciprofloxacin at 100 times the concentration required to inhibit cell growth, known as the minimum inhibitory concentration (MIC, 10 µg/ml) [10]. Consequently, all cells initially exposed to high ciprofloxacin concentrations either fail to grow and/or are lost from the substrate (**Fig. 4Aiii**) to be rapidly flushed away. As expected however, some surface-attached cells on the antibiotic-free side of the gradient begin to migrate up the ciprofloxacin gradient (**Fig. 4Aiv**). To follow this migration, we image conduits for up to 36 h post-inoculation. After ∼24 h, migrating WT cells accumulate in regions containing high ciprofloxacin concentrations (**Fig. 4B**); in contrast, *ΔpilG* mutants remain largely restricted to the media-only side of the conduit. Analysis of the trajectories of more than 1,000 WT cells (see **Materials and Methods**) shows that in the taxis zone, there were 2-fold more trajectories moving towards the antibiotic compared to those moving away from it (shown in blue and orange, respectively, in **Fig. 4Di**) – a bias confirmed in a rose plot (**Fig. 4Dii**). We also use trajectories to detect when cells abruptly reverse the direction of their movement. It has been shown previously that these cells generate chemotaxis by deploying these “reversals” more frequently when moving away from, rather than towards chemoattractants [21], [24], and this is confirmed here (**Fig. 4Diii**).

#### Reconfiguring circuits on a microscope to isolate migrating cells

After ∼24 h, many cells have undergone chemotaxis towards ciprofloxacin and so we next isolate this migrating sub-population from those establishing a biofilm in antibiotic-free regions of the circuit. As we have seen, this can only be achieved with continuous flow [10]. Therefore, we use the 3-D printed adaptors discussed above to attach a PTFE stylus to the microscope condenser (**Fig. 2Bii**), and then use the motorized stage on the microscope to move the stylus relative to the dish. This allows us to build a new fluid wall down the length of the central conduit separating the bulk of the migrating population from the antibiotic naïve population of cells (**Fig. 4C, Fig. S4**; **Materials and Methods**). Once this wall is built, flows through the circuit are stopped and two additional walls were built across the width of the central conduit above and below the taxis zone to completely isolate those cells that have undergone chemotaxis in an isolated fluid chamber containing ∼100 nL (**Fig. S4C**).

#### Extracting chemotaxing cells and assessing their viability

As discussed in the companion paper [22], a key question is whether migrating cells in the experiment develop antibiotic resistance and therefore retain long-term viability. To assess long-term viability using our method, circuits can be transferred back to the printer and the contents of the isolation chamber extracted (**Fig. S5A**). This fluid (containing any planktonic cells) is then plated onto agar plates containing antibiotic-free growth medium to monitor colony growth. The viability of cells remaining within the isolation chamber is also monitored continuously. Any residual ciprofloxacin in the chamber is removed and replaced with antibiotic-free media, dishes returned to the microscope, and the entire isolation chamber imaged for at least 36 h to detect any cell growth and division. This experiment revealed that neither cells extracted from the isolation chamber, nor those remaining behind in it, retained long-term viability. Why then does *P. aeruginosa* move towards antibiotics? It appears that chemotaxis towards antibiotics may be driven by an evolved response of surface-attached *P. aeruginosa* to move towards toxins produced by neighbouring competitors in an attempted counter-attack [22]. However, in the case of antibiotic gradients, this counter-attack proves futile, and the attacking cells die.

## Discussion

We report two general methods for isolating desired sub-populations of cells from within microfluidic circuits that are formed by fluid walls (**Fig. 1**) and exemplify them by isolating highly-enriched chemotaxing macrophages or bacteria relative to their non-responding counterparts. In the first approach, circuits are fabricated, manipulated, and reconfigured on a custom-built three-way traverse. As these operations are performed on the traverse without live microscopy, we design 3-D printed components that fit on condensers of conventional inverted microscopes to transform them into micromanipulators to reconfigure circuits (**Fig. 2**). This allows specific sub-populations of cells to be isolated using high-precision real-time microscopy as a guide. Our approach also allows flow through circuits to continue uninterrupted as critical isolating walls are built. Using the traverse, we first isolate macrophages chemotaxing towards C5a (**Fig. 3**). We then use the 3-D printed adaptors to target and isolate bacteria migrating towards antibiotic in real-time on a microscope (**Fig. 4**). Note that we also isolate macrophages using a similar technique and a different type of inverted microscope (**Fig. S8**).

Uniquely, our approach here allows isolation of desired sub-populations of adherent migratory prokaryotic and eukaryotic cells as flow and imaging continue. In both cases, the isolated sub-population can be extracted from the circuit and analysed using a range of different techniques. For example, RNA sequencing could be applied to identify which transcripts increase in number in a chemotactic sub-population. Moreover, as techniques exist to build fluid walls with sufficient accuracy to isolate single cells [4],[12], the methodology outlined here could even be applied with single-cell transcriptomics. In this way, our method greatly increases the range of analytical methods that can be combined with microfluidic experiments.

## Materials and Methods

### Strains, media and culture conditions

To obtain BMDMs, bone marrow was extracted from hind-limbs of hCD68-GFP transgenic mice bred on a C57BL/6 background – C57BL/ 6-Tg (CD68-EGFP) 1Drg/ J (026827, The Jackson Laboratory). Bone marrow cells were cultured (7 d, 37°C, 5% CO_2_) in high-glucose Dulbecco’s Modified Eagle’s Medium (DMEM, Sigma) enriched with 10% L929-conditioned media (containing macrophage colony-stimulating factor), 10% foetal bovine serum (FBS, Sigma) and 1% penicillin plus streptomycin (P/S, Gibco). 8 mL growth media was used for the first 4 d; then, 3 mL media was removed and replaced with 5 mL fresh media. Cells were plated into circuits on day 6, and chemoattractants added on day 7. Chemoattractant used in BMDM experiments was recombinant mouse complement component 5a (C5a, 10 nM; R&D Systems).

The wild-type *P. aeruginosa* strain used throughout this study was PAO1 (Kolter collection ZK2019) expressing yellow fluorescent protein (YFP) from a constitutive promoter [25]. As an in-experiment control, we used a PAO1 strain harbouring an in-frame deletion of *PilG* (*ΔpilG*, [10]). All bacteria were grown from frozen stocks overnight in LB medium (ThermoFisher; 37°C; 250 rpm) and sub-cultured (1:30 dilution) in tryptone broth (TB, 10 g L^-1^ Bacto Tryptone; ThermoFisher) for 2.5 h (37°C; 250 rpm). Cells were then diluted in TB to an optical density (OD) at 600 nm of 0.5 before inoculation into circuits.

### Imaging

BMDMs were imaged at a rate of 3 frames h^-1^ using an IncuCyte ZOOM, a commercially available live-cell imaging system (Sartorius, Gottingen, Germany). Bright-field images of *P. aeruginosa* were captured at a rate of 1 frame min^-1^ using a Zeiss Axio Observer inverted microscope with a Zeiss AxioCam MRm camera, a Zeiss Definite Focus system and a 20X Plan Apochromat air objective. Experiments that imaged YFP-labelled cells used a Zeiss HXP 120 illuminator with an exposure time of 150 ms. In all experiments, cell movement was followed in real-time using bright-field images that were processed using Fiji [26] and analysed using Matlab as described previously [21].

### Circuit design and fabrication

To study chemotaxis, we used ‘m’-shaped microfluidic circuits to study both BMDMs and surface-attached *P. aeruginosa*, although the latter circuits were smaller in size (**Fig. S2**; note that BMDM cells are ∼20 µm in diameter, and *P. aeruginosa* cells a few microns in length). All needles, glass syringes, and Teflon tubing were internally sterilised and washed by infusing 70% ethanol followed by sterile culture medium (either DMEM or TB), and then externally sterilised by bathing in 70% ethanol for 1 min followed by sterile medium for 1 min.

#### BMDM circuit

The BMDM circuit was printed using a modified, custom-made printer (iotaSciences Ltd, Oxford, UK). The three-axis traverse on the printer holds a 25 G dispensing needle (Adhesive Dispensing Ltd) connected by PTFE tubing (Cole-Parmer) to a 250 µL glass syringe (Hamilton) controlled by a syringe pump (iotaSciences Ltd). The dispensing needle was brought 300 µm above the surface of the dish and then infused (10 µL/min) DMEM + 10% FBS + 10% L929 + 1% P/S + 2% fibronectin as it drew the circuit [3]. Once completed the circuit was overlaid with 2 mL FC40 (iotaSciences Ltd) to prevent evaporation. Circuits were printed on both 60 mm tissue culture-treated dishes (Corning, 430166), and 60 mm suspension dishes (Corning, 430589). As BMDMs are extremely adherent, suspension dishes were used when cells were to be extracted (**Fig. S3**).

#### P. aeruginosa circuit

The bacterial circuit was printed on the surface of untreated 50 mm glass-bottomed dishes (MatTek; the diameter of the glass bottom is 30 mm) as described above except that the needle was lowered to within 200 µm of the glass surface, and then moved across the dish as it infused (10 µL/min) DMEM + 10% FBS. After printing, circuits were immediately overlaid with 2 mL FC40. A thin and straight FC40 wall (length ∼1 mm, width ∼100 µm) was then built at the junction where the two-inlet arms merge into the central conduit (**Fig. S2B**), to allow incoming streams to meet roughly parallel to one another. This wall was made by jetting (**Fig. S6**; [12]). Briefly, the dispensing needle was used to generate a jet of FC40. When this jet is targeted onto an existing circuit, the FC40 jet forces aside the medium making up the circuit, analogous to the PTFE stylus driving medium aside. The FC40 subsequently remains stuck to the dish, held in place by interfacial forces. We therefore jetted FC40 (6.5 µL s^-1^) through a 70 µm diameter needle (Oxford Lasers) positioned 0.5 mm above the aqueous circuit as the needle traversed in a straight line (1000 mm/min) to create the ∼100 µm wide dividing wall.

### Exposing BMDMs to a gradient of C5a

Following circuit fabrication, the sink was sealed off from the central conduit by building a new wall across the width of the conduit with a PTFE stylus. This allowed cells to be added directly within the conduit, without flowing into the sink due to differences in pressure across these two reservoirs – the larger size of the sink (and hence radius of curvature) results in a lower Laplace pressure [3] and therefore any solution added to the conduit will flow directly into the sink down the resulting pressure gradient. Note that this was not necessary when carrying out experiments with *P. aeruginosa* because we were able to use a much higher inoculum density, allowing many cells to attach before drifting into the sink.

BMDMs (5 µL with 1000 cells µL^-1^) were infused into the conduit, and the dish incubated overnight to allow cell attachment. Using a hydrophilic stainless-steel 25 G needle (identical to the one used to print circuits), the sink was reconnected with the conduit by manually dragging the needle between the two interfaces and allowing fluid walls to merge back with one another. Three syringes (Hamilton) were then loaded onto syringe pumps (PhD Ultra, Harvard Apparatus): one 5 mL glass syringe filled with DMEM + 10% FBS + 10% L929 + 1% P/S, another identical one with the same media + 10 nM C5a, and a third 10 mL glass syringe with 1 mL growth media. The 5 mL syringes were mounted on the same syringe pump, while the 10 mL syringe was fitted on a separate syringe pump. All three syringes were connected to 25 G needles (Adhesive Dispensing Ltd) via PTFE tubing (1 m). A 3-D printed needle holder that attaches to the outside edge of the dish then aids manual placement of needles in the circuit (**Fig. S7**). To ensure needles were inserted 100 µm above the glass surface, the holder was first placed on a separate dish containing a glass coverslip (Menzel-Glaser #1 thickness) glued to the bottom. Needles were then lowered through the holder until they came into contact with the coverslip, at which point the needles were bent over the holder to ensure they remained at the same height throughout an experiment (**Fig. S7B,iii**). With needles in the holder at an appropriate height, the holder was transferred to the dish containing the circuit. The 5 mL syringes ± chemoattractant were connected to the two arms of the circuit, and the 10 mL syringe to the sink. The circuit was then overlaid with an additional 4 mL of FC40 and placed in the IncuCyte (inside an incubator at 37°C, 5% CO_2_). The 5 mL syringes were set to infuse at 0.5 µL min^-1^, while the 10 mL syringe was set to extract at 1 µL min^-1^.

### Exposing *P. aeruginosa* to a ciprofloxacin gradient

After fabrication, circuits were infused (0.5 µL min^-1^) with 1.5 µL TB through each of the two inlet arms to flush out the DMEM + FBS used during fabrication. Flushed media flowed into the sink at the end of the central conduit and was subsequently removed. *P. aeruginosa* cells (3 µL with OD at 600 nm of 0.6) were then infused (0.4 µL min^-1^) into one arm of the circuit upstream of the junction, and the circuit was left for 10 min to allow cells to attach to the glass surface.

Two 500 µL glass syringes (Hamilton) were then used to set up the antibiotic gradient – one containing TB plus Chicago Sky Blue dye (0.05 mg mL^-1^) to enable visualisation of the (reverse) concentration gradient, the other with TB plus 10 µg mL^-1^ ciprofloxacin (corresponding to 100 times the MIC). This dye does not influence cell motility in surface-attached cells [21]. A third plastic 10 mL syringe (Becton Dickinson Plastipak) was loaded with 1 mL TB plus 10 µg mL^-1^ ciprofloxacin (corresponding to 100 times the MIC) and a fourth plastic 1 mL syringe (Becton Dickinson Plastipak) with 1 ml TB plus 0.3 µg mL^-1^ ciprofloxacin (corresponding to 3 times the MIC) and Chicago Sky Blue dye (0.05 mg mL^-1^). Each syringe was fitted into an individual syringe pump (PhD Ultra, Harvard Apparatus), and connected to a 25 G needle (Adhesive Dispensing Ltd) via PTFE tubing (1 m). The four needles were then placed into a needle holder (**Fig. S7**) and set to the appropriate height using the same ‘zeroing’ technique as before. The holder was transferred to the dish containing the circuit, and needles lowered into the circuit: the needle connected to the glass syringe plus TB was lowered into one arm, the second connected to the glass syringe plus antibiotic was lowered into the other arm, the third connected to the 10 mL plastic syringe was lowered into the sink, and the fourth connected to the 1 mL plastic syringe was lowered into the antibiotic-containing arm upstream of the other inserted needle (**Fig. S7A**). A further 4 mL FC40 was then overlaid on the circuit to prevent evaporation during the experiment.

To generate a concentration gradient of ciprofloxacin, TB plus 10 µg mL^-1^ ciprofloxacin was infused into one inlet arm, and TB plus dye into the other (0.1 µL min^-1^ each). The two fluids then joined as laminar streams after the dividing wall and flowed through the central conduit into the sink. Diffusion between streams then generated two concentration gradients across the conduit width; one due to antibiotic, the other due to dye. Visual observation of the dye gradient allowed us to be sure that flows were stable and laminar, and that diffusion gradients were established. Fluid was also simultaneously withdrawn at an equivalent flow rate (0.2 µL min^-1^) through the needle in the sink into the 10 mL plastic syringe. After ∼12 h, rates of infusion and withdrawal were halved to widen the ciprofloxacin gradient and so increase the number of cells that were exposed to the antibiotic gradient. All antibiotic-taxis experiments were performed at 22°C using a custom-designed microscope incubation chamber with both heating and cooling modes (Digital Pixel Cell Viability and Microscopy Solutions, Brighton, UK).

### Determining maximum gradient widths

Fluid walls change shape in response to changes in pressure and in a straight conduit, the height will decrease from the point of highest pressure (i.e., where the needle infuses media) to the point of lowest pressure (i.e., the conduit exit near the sink). It was recently shown that conduit heights and flow velocities in fluid-walled circuits can accurately be predicted from a simple semi-analytical model [27]. Using this equation, we determined the average maximum height (*h*_*max,avg*_) and particle velocity (*u*_*max,avg*_) for each conduit in this study in order to estimate the average amount of time a particle spends in the conduit as 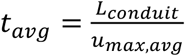, where *L*_*conduit*_ corresponds to the length of the conduit observed. We then used this time to calculate the characteristic distance the chemoattractant diffuses across the width of a conduit (*x*_*particle*_) using the characteristic length scale of diffusion 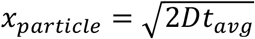, where *D* is the diffusion coefficient of the chemoattractant.

We next determined the diffusion coefficient of C5a, *D*_*c*5*a*_. Using the Stokes-Einstein equation for the diffusion of a spherical particle in water of radius *r* as 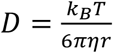 (*k*_*B*_ = Boltzmann’s constant, *T* = absolute temperature, *η* = dynamic viscosity), the Stokes radius of a C5a molecule is estimated using its molecular weight, *MW*_*c*5*a*_ = 9 kDa, as *r* = 1.82 nm. Then, using *η*_*water*_ = 0.7 cP and *T* = 310.15 K (37°C), we find *D*_*c*5*a*_ = 1.8 × 10^−10^ m^2^/s. Hence, for the BMDM circuit, *Q* = 1 µL min^-1^ (*Q* = flow rate), *h*_*max,avg*_ ≈ 100 µm, *u*_*max,avg*_ ≈ 10,800 µm/min, *L*_*conduit*_ = 3 mm, therefore the maximum gradient width obtained within the conduit is *x*_*c*5*a*_ ≈ 80 µm. For the *P. aeruginosa* circuit, *Q* = 0.2 µL min^-1^, *h*_*max,avg*_ ≈ 40 µm, *u*_*max,avg*_ ≈ 21,600 µm min^-1^, *L*_*conduit*_ = 2 mm, *D*_*cipro*_ = 4 × 10^−10^ m^2^ s^-1^ [28], therefore, the maximum gradient width obtained within the conduit is *x*_*cipro*_ ≈ 70 µm.

### 3-D printed adaptors for easy manipulation of circuits on and off the microscope

The workflow requires transfer of circuits between microscope stage and printer. To ensure circuits retain the same orientation on both platforms, dishes were fitted into a carrier bearing a single triangular protrusion (positioned 63° counter-clockwise from the top). This carrier was designed to fit in a fixed orientation into holders built into both the printer and microscope stages that bore complementary triangular indents (**Fig. 2**). Both carrier and holder were designed using Onshape (Boston, MA) and fabricated using a Flashforge Finder 3-D printer (Zhejiang, China). Carriers were designed for both Corning and MatTek dishes and can be easily adapted to fit a dish from any manufacturer.

### Calibrating a circuit before reconfiguration on a microscope

We designed a simple adaptor to fit a PTFE stylus onto the condenser of conventional microscopes, allowing us to reshape circuits without interrupting flows (**Fig. 2**). The adaptor was designed using Onshape and fabricated using a Flashforge Finder 3-D printer (**Supplementary STEP files**). To reconfigure circuits, the stylus was first held to one side of the condenser turret to not interfere with imaging. We then calculated the distance between the centre of the objective lens (the centre of the field of view) and the position of the stylus. To perform this calibration, we designed an ‘L’-shaped conduit placed above the main circuit (**Fig. S4A,B**). The stylus was then lowered onto the surface of the dish in the gap between both arms of this ‘L’ and the coordinates of this position (*p1*) were recorded. Next, we moved the microscope stage vertically, moving the stylus perpendicularly through the horizontal arm of the ‘L’ (‘path 1’ in **Fig. S4B**). After returning the microscope stage to *p1*, we moved the stage horizontally, thus moving the stylus perpendicularly through the vertical arm of the ‘L’ (‘path 2’ in **Fig. S4B**). This created both a horizontal and vertical wall through the arms of the ‘L’-shaped conduit (**Fig. S4B**). The offset between the coordinates of *p1* and the centre of these vertical and horizontal walls corresponded to the *x*- and *y*-offsets respectively between the objective lens (*p2*) and attached stylus. Using these offsets then allowed us to position the stylus precisely within any observable region of the dish. The precision with which any new walls were built was thus determined by the movement resolution of the microscope stage (typically in the micrometre range) and the width of the stylus (typically tens of micrometres), although this precision can be slightly reduced by play in the ring that holds the stylus.

### Isolating and extracting chemotaxing BMDMs both on and off the microscope

After ∼28 h in the circuit, BMDMs began migrating towards higher C5a concentrations, creating a region of high cell density (**Fig. 3, Fig. S3, Fig. S8**); the first fluid wall was built to isolate this band from cells remaining in low C5a concentrations. Since BMDMs are so adherent [19], flows can be stopped prior to reconfiguring the circuit without risking cross-contamination from detaching cells. The reconfiguration was then performed by placing the dish on the printer, which builds the new walls. Wall coordinates were determined by analysing cell trajectories, and the stylus was washed after building each wall with 70% ethanol to prevent cross-contamination (**Fig. 3, Fig. S3**). Additionally, circuits were also reconfigured on the stage of an Olympus IX53 inverted microscope using a modified version of the adapter holding the PTFE stylus on the condenser. The stylus was positioned directly above the objective lens and the circuit was dragged past it in real time by moving the stage of the microscope as described previously (**Fig. S8**).

#### Automated retrieval of isolated BMDMs

The circuit was returned to the printer to extract the small volume remaining in the isolation chamber. First, the chamber was washed twice by sequentially adding and removing 0.5 µL PBS (5 µL min^-1^), followed by 1 µL 10 mM EDTA (a metal chelator that reduces adhesion of integrins [19]) in PBS (5 µL min^-1^), and then incubating for 10 min at 37°C. The chamber was then gently but firmly tapped from below the dish using a pen to ensure that most cells were dislodged (confirmed visually on the microscope). To extract dislodged cells, a volume of 1.5 µL was withdrawn (5 µL min^-1^) from the chamber using an extracting needle that had been pre-loaded with 20 µL cell media to ensure that withdrawn cells were immediately exposed to nutrient-rich media and to neutralise EDTA (**Fig. S3**). The needle then dispensed (100 µL min^-1^) 5 µL of its contents into an Eppendorf tube containing 95 µL cell media, the contents of which were then transferred to a 35 mm TC-treated dish (Corning, 430165) and incubated for 24 h to assess cell viability.

#### Assessing BMDM viability post-extraction

To assess viability of extracted BMDMs, we first counted the number of cells contained within the isolation chamber immediately after building the surrounding walls and compared this to the number of cells extracted and transferred to the new dish. We then defined a specific region of interest including only the population of chemotaxing cells (ROI – dashed outline in **Fig. S3B-D**) from which we extracted ∼87% of cells. In total, ∼80% of these extracted cells re-attached to the surface of the dish and displayed their characteristic pseudopodia, indicating they remained viable (**Fig. S3F**).

### Isolating and extracting migrating *P. aeruginosa* cells

We first sought to build a fluid wall along the length of the central conduit to isolate specific sub-populations of migrating bacteria from the high density of non-migrating cells present in antibiotic-free regions (**Fig. 4C**). As bacteria migrate up the antibiotic gradient, they experience a steady increase in antibiotic concentration. However, building a separating fluid wall along the centre-line of the central conduit to isolate migrating cells from the bulk population would disrupt the ciprofloxacin gradient and rapidly expose all migrating cells to the maximum ciprofloxacin concentration (100 times the MIC). To avoid exposing cells to such a dramatic increase in concentration, we first decreased the concentration of antibiotic perfused to 3 times the MIC. We thus prepared circuits for reconfiguration by additionally infusing (0.05 μL min^-1^) TB plus ciprofloxacin at 3 times the MIC through the same circuit arm in which we infuse TB plus ciprofloxacin at 100 times the MIC. This temporarily doubles the flow rate through this circuit arm, which shifts the gradient towards the antibiotic-free side. Immediately after obtaining visual confirmation of this shift by observing a change in the position of the dye gradient (∼20 s), infusion through the syringes containing antibiotic-free TB and the syringe containing TB plus ciprofloxacin at 100 times the MIC was paused in order to fill the conduit with ciprofloxacin at 3 times the MIC.

At this stage, we built an initial fluid wall along the length of the central conduit as shown in **Fig. 4C**. After building this first wall, we continued to infuse TB plus ciprofloxacin at 3 times the MIC for 5 min to wash away any antibiotic-naïve planktonic cells that could have been inadvertently displaced from antibiotic-free regions of the circuit during circuit reconfiguration. Flow was then halted and two additional walls were built across the width of the central conduit to completely seal off an isolated fluid chamber containing migrating bacteria (**Fig. S4C**). Before building each wall, the stylus was raised and sterilised with 70% ethanol to avoid contamination from cells it may have contacted previously.

#### Extracting planktonic P. aeruginosa from the isolation chamber

Isolated bacteria remained in fluid chambers containing ciprofloxacin at 3 times the MIC for 2 h in order to kill any antibiotic-naïve cells [10]. Cells were monitored by imaging (0.1 frame min^-1^) and whilst most cells remained surface-attached, some detached and were lost from the field of view. We therefore analysed the viability of both surface-attached and planktonic populations. We analysed the planktonic fraction by extracting chamber contents and plating them onto antibiotic-free LB agar plates. To this end, culture dishes were returned to the printer, and an extracting needle was pre-loaded with 50 µL TB media plus Chicago Sky Blue dye (0.05 g mL^-1^); this ensured that withdrawn cells were immediately exposed to a nutrient-rich environment, whilst the blue dye allowed us to visually monitor the exchange of the nanolitre scale volumes in the chamber. The needle was lowered to within 100 µm of the dish surface, and 1 µL was withdrawn (2 µL min^-1^). This volume exceeded the total amount of media in the chamber, so some FC40 was also withdrawn. We then dispensed 5 µL from the needle at a flow rate of 100 µL min^-1^ into an Eppendorf tube containing 95 µL LB medium to ensure that any bacteria that had been withdrawn into the needle were likely flushed out. The contents of the Eppendorf tube were then spread on LB agar plates, incubated (92 h, 37°C) and colony growth monitored.

#### Assaying viability of surface-attached P. aeruginosa

To analyse the viability of cells remaining attached to the surface of the dishes within the isolated fluid chamber, the chamber (formed of TB overlaid with FC40) was washed five times with TB media plus dye to dilute the antibiotic to a negligible level. The contents of the chamber during each wash were also recovered in separate Eppendorf tubes containing 95 µL LB media, and later plated to monitor colony growth. Each wash involved infusing (0.1 µL min^-1^) 0.1 µL TB media plus dye into the chamber and removing 1 µL from it (including some FC40). The dishes were then returned to the microscope and the bottom-surface of the entire chamber imaged for 60 h (0.1 frames min^-1^) to monitor cell movement and viability. After 60 h, we infused (at a flow rate of 0.1 µL min^-1^) 0.1 µL of exponential-phase cell suspension (diluted to an OD at 600 nm of 0.015) into the chamber to introduce ∼25 viable cells. These cells grew and rapidly filled the entire chamber after ∼12 h; this acted as a positive control indicating that the concentration of antibiotic remaining in the isolated chambers was negligible (**Fig. S5B**).

### Image analysis

To quantify the motility of individual bacteria and macrophages, we tracked thousands of cells using a cell-tracking approach we developed previously [21]. Briefly, a series of bright-field images were stabilised and the background pixel intensity normalised using the ‘Image Stabiliser’ and ‘Normalise Local Contrast’ plugins in Fiji [26]. The background was then subtracted (using ‘Subtract Background,’ in Fiji) and the pixel intensity inverted to generate an image series with high-pixel intensity cells on a low-pixel intensity background. These cells were then tracked using the open source ‘Trackmate’ plugin for Fiji [29]. Further analyses of the resulting trajectories including the quantification of reversal rates and cells’ chemotactic bias were performed in Matlab, as previously described [21].

## Supporting information

CAD files

## Acknowledgments

This work was supported by iotaSciences Ltd. (who provided a scholarship for C. D.), the Biotechnology and Biological Sciences Research Council (BB/R018383/1 – J. H. R. W.), the Impact Acceleration Account of the Biotechnology and Biological Sciences Research Council (P. R. C. and E. J. W.), awards from the Medical Research Council under the Confidence in Concept scheme (MC_PC_15029 to P. R. C. and E. J. W), the British Heart Foundation graduate studentship (FS/17/68/33478 to A. N. R.), the Human Frontier Science Program (RGY0080/2021), EPSRC Pump Priming Award (EP/M027430/1) and BBSRC New Investigator Grant (BB/R018383/1) to W. M. D., and the European Research Council Grant (787932) and Wellcome Trust Investigator award (209397/Z/17/Z) to K. R. F.

## Author Contributions

C. D. and J. H. R. W. contributed equally. C. D., J. H. R. W., P. R. C., W. M. D., K. R. F. and E. J. W. designed the research; C. D. and J. H. R. W. performed experiments and analyzed data; A. N. R. maintained and provided BMDMs; and all authors worked on the paper.

