## Supplementary Information for "Reconfigurable microfluidic circuits for isolating and retrieving cells of interest"

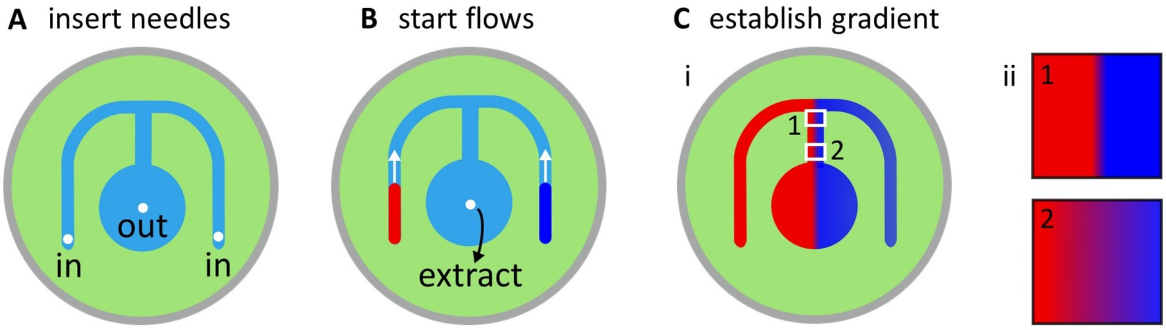


**Figure S1. Establishing gradients in fluid-walled circuits in an 'm'-shaped circuit. (A)** Needles connected to syringe pumps via tubing are inserted into the arms and sink. **(B)** Red and blue dyes are pumped into the two arms and withdrawn from the sink at an equivalent flow rate. **(C)** As laminar streams of red and blue dyes meet in the central conduit, they flow side by side and diffuse laterally across the conduit. **(i)** White boxes labelled 1 and 2 illustrate regions with differences in diffusional mixing along the length of the conduit, and correspond to insets **(ii,1)** (where the gradient is steeper near the top of the conduit) and **(ii,2)** (where the gradient widens along the bottom of the conduit as particles have more time to diffuse), respectively.

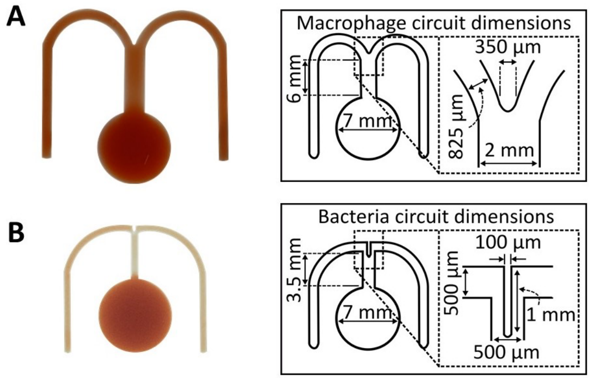


**Figure S2. Dimensions of circuits used to study macrophage and bacterial chemotaxis. (A)** Left: micrograph of the circuit used with macrophages (red dye added to aid visualisation). Right: circuit dimensions. **(B)** Left: micrograph of the circuit used with bacteria (red dye added to aid visualisation). Right: circuit dimensions.

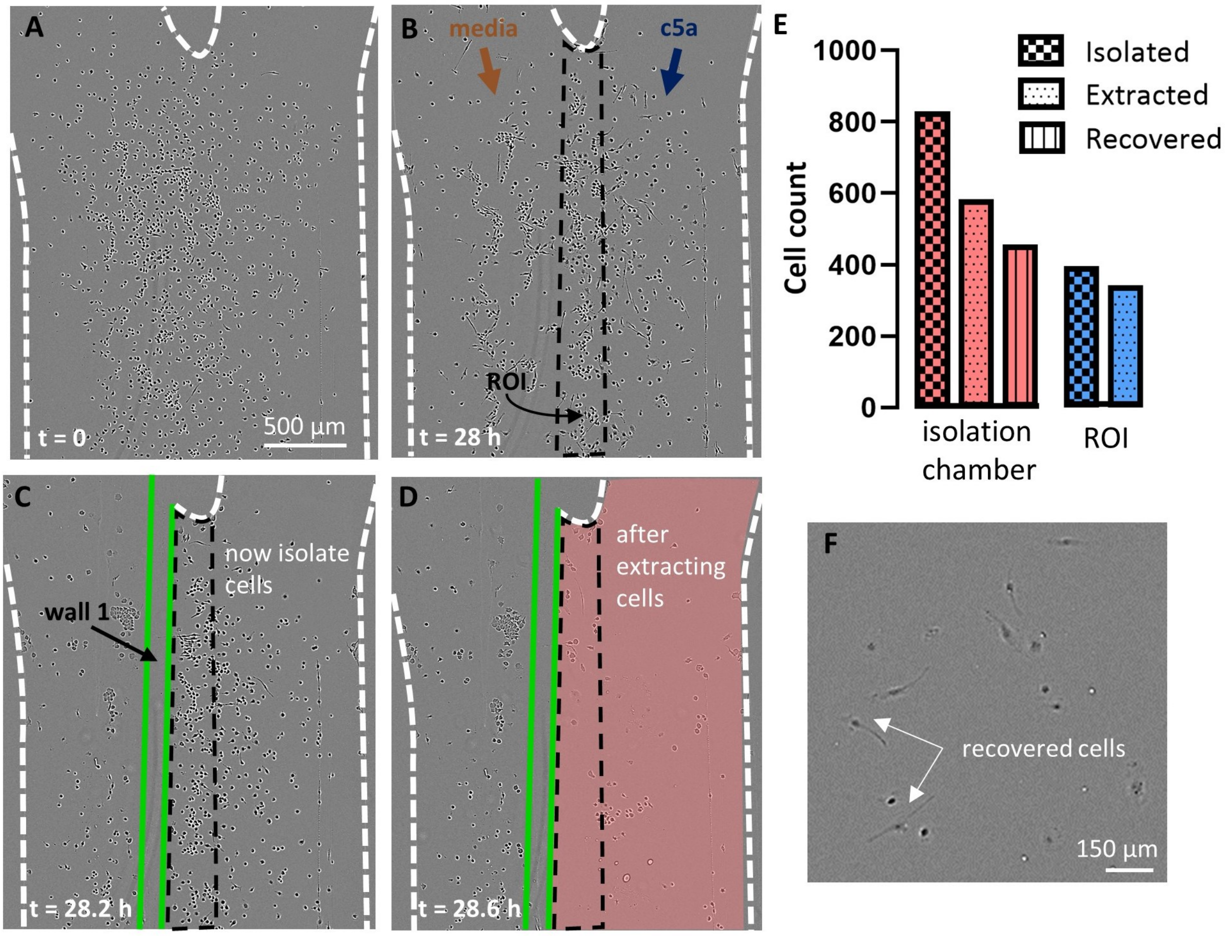


**Figure S3. Isolation, extraction, and recovery of chemotaxing BMDMs.** Phase-contrast images were collected on the IncuCyte. **(A)** Image of BMDMs taken immediately after plating in the central conduit of the 'm'-shaped circuit. **(B)** Image of the conduit after 28 h as laminar streams of media and C5a flow over cells. Some cells sense the C5a gradient and migrate to the right to leave a zone with fewer cells. Analysis of the movie allows us to determine the region containing migrating cells (region of interest 'ROI', outlined). **(C)** Image of newly-built 'wall 1' (green, made using the stylus on the printer). A further two walls (not shown in image) are then built to completely seal off the region rich in chemotaxing cells from the rest of the circuit. **(D)** Image of the isolation chamber (pink) containing the ROI after extracting cells using the printer. **(E)** Graph quantifying the number of cells isolated, extracted, and recovered from within the isolation chamber and the ROI. Within the isolation chamber (which includes the ROI), 70.2% of isolated cells were successfully extracted with the printer (which includes 87% of the cells in the ROI). 80% of these extracted cells were later recovered after plating onto new culture dishes. **(F)** Image of recovered BMDMs from the isolation chamber after transfer into new dishes and incubation for 24 h. Cells have reattached and present their characteristic pseudopodia.

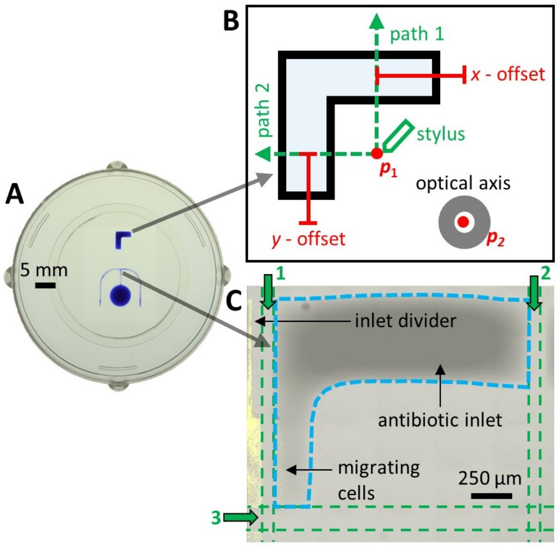


**Figure S4. Reconfiguring microfluidic circuits on a microscope to isolate chemotaxing *P. aeruginosa* without interrupting flow. (A)** Image of dish with 'm' and 'L'-shaped circuits (both contain blue dye to aid visualisation). **(B)** Plan of 'L'-shaped circuit used to ascertain *x*, *y* offsets of stylus from the optical axis. After printing circuits, the condenser is lowered so the attached stylus touches the surface of the dish at an arbitrarily-chosen point ($p_1$) lying orthogonal to arms of the 'L'. Next, the motorized stage moves so that the stylus follows paths 1 and 2 over the dish surface. During movement, media in contact with the glass is displaced by the stylus, and – as FC40 wets glass better than media – FC40 walls pinned to the glass are built along paths 1 and 2. Mid-points in these newly-built walls are imaged through the objective lens, and *x*, *y* offsets from the optical axis ($p_2$) recorded. **(C)** Overlaid fluorescence and bright-field images of 'm'-shaped circuit (at junction between antibiotic arm and central conduit) that contains wild-type YFP-labelled *P. aeruginosa* cells, (a high density can be seen as the yellow on the left). Live-cell imaging of the biofilm over the previous ~24 h shows that many cells to the right of what will become path 1 were chemotaxing towards ciprofloxacin. Therefore, new FC40 walls were built successively along paths 1-3 (green arrows) to isolate these cells from others (see **Materials and Methods**). The volume within the dashed blue outline is now isolated from the rest of the dish. In this low-power view, individual cells yield insufficient fluorescence signal to be detected.

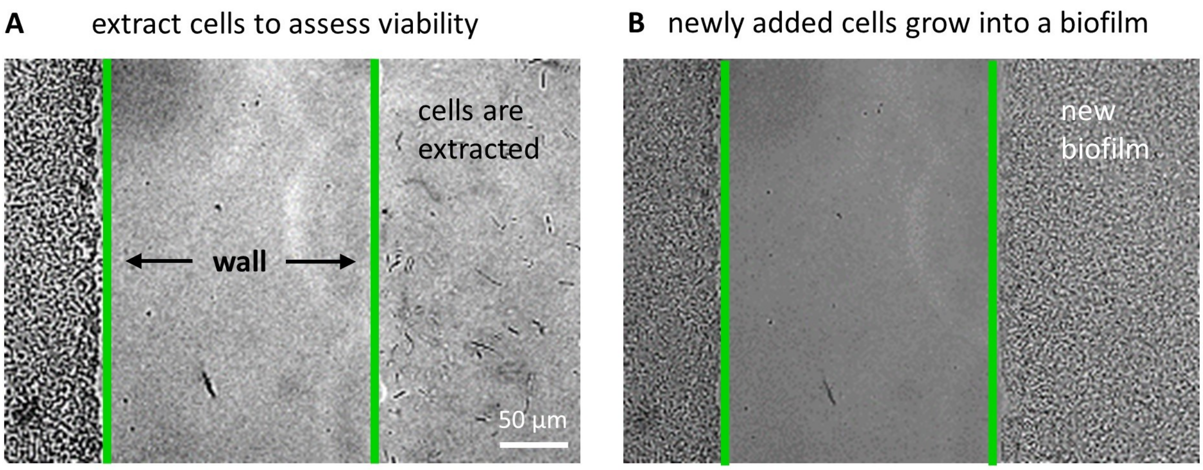


**Figure S5. Phase-contrast images showing that *P. aeruginosa* in newly-reconfigured circuits can be extracted or added to monitor division potential. (A)** Image after extracting cells to the right of the wall; few cells remain in the chamber, and the dense biofilm to the left serves as a control. Subsequent plating of extracted cells on LB plates yields no colonies, showing none can divide. **(B)** Image taken 12 h after re-adding ~25 new WT cells in antibiotic-free media to the extracted area shows that washing the chamber thoroughly can remove essentially all antibiotic. The chamber is now completely covered in a biofilm of WT cells (indicating that cell growth and division is possible in the isolated region and that there is negligible residual antibiotic).

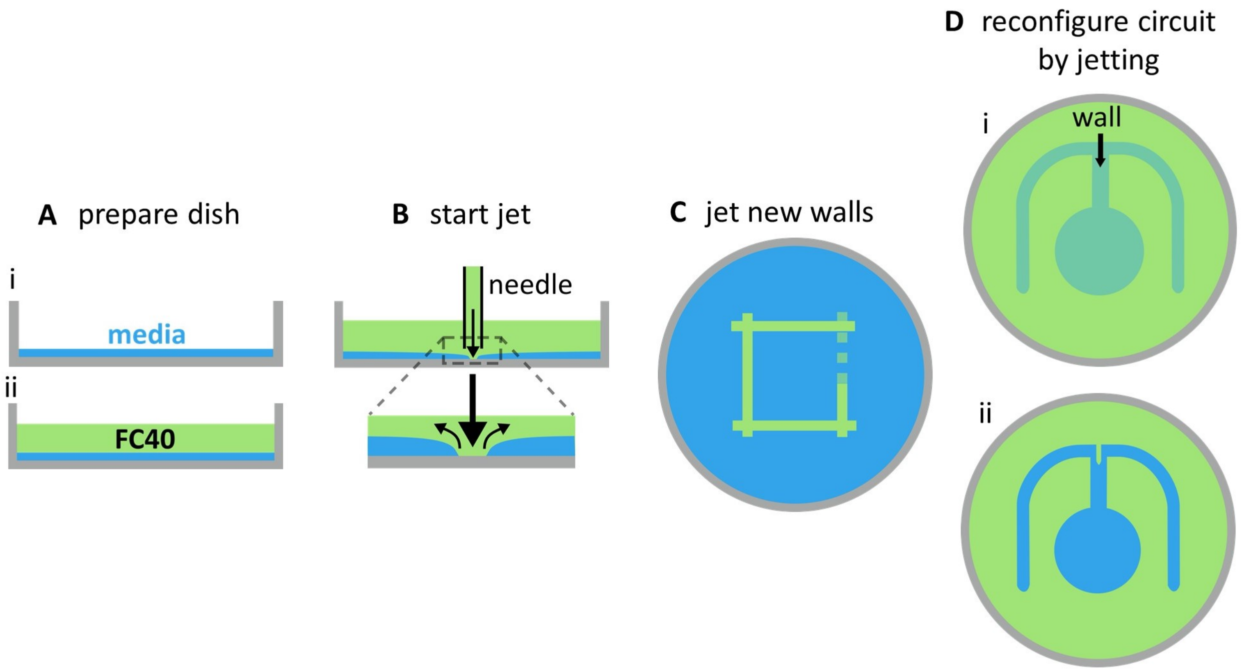


**Figure S6. Jetting enables fine-scale reconfiguration of microfluidic arrangements. (A)** Preparing a dish for jetting. **(i)** A virgin Petri dish is overlaid with a thin film of media. **(ii)** Media is overlaid with a layer of FC40. **(B)** A dispensing needle loaded with FC40 is held in the FC40 above the dish. The needle 'jets' FC40, forcing media aside and allowing FC40 to wet the dish. FC40 wets the surface better than media, so it remains stuck to it. **(C)** Fluid walls may be built by moving the needle over the dish, shown here in the fabrication of a square. **(D)** We use jetting to create just the thin dividing wall at the point where both arms of the 'm'-shaped circuit used for *P. aeruginosa* meet (**Fig. S2B**). **(i)** The jetting needle moves along the path indicated by the black arrow to build a new wall. **(ii)** Representation of the completed circuit after jetting.

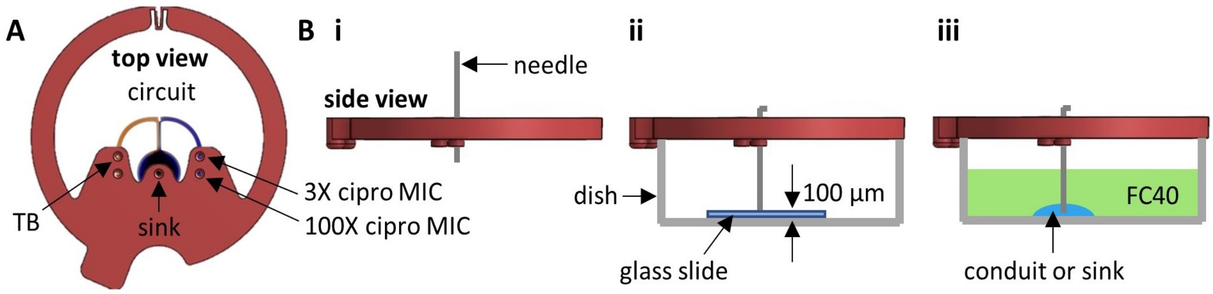


**Figure S7. Calibrating the height of needles used to infuse and withdraw fluid through fluid walls. (A)** Image (top view) of a needle holder (red, CAD model) attached to a dish containing an 'm'-shaped circuit into which red and blue dyes have been pumped into the side arms (the edge of the dish is hidden by the circumference of the needle holder. The holder has holes (labelled) through which needles can be inserted that are positioned directly above appropriate points in the circuits. **(B)** Adjusting the height of an input needle (CAD views from side -/+ cartoons below; a similar procedure is used with the other two needles). **(i)** The needle is inserted through the needle holder. **(ii)** The needle is fitted to the dish with a glass coverslip (height 100 µm) glued to the bottom, and lowered until it contacts the coverslip; then, the needle is bent to fix its vertical position in the holder. **(iii)** When the needle holder is placed on a dish with a circuit, the needle is now at the correct height relative to the circuit.

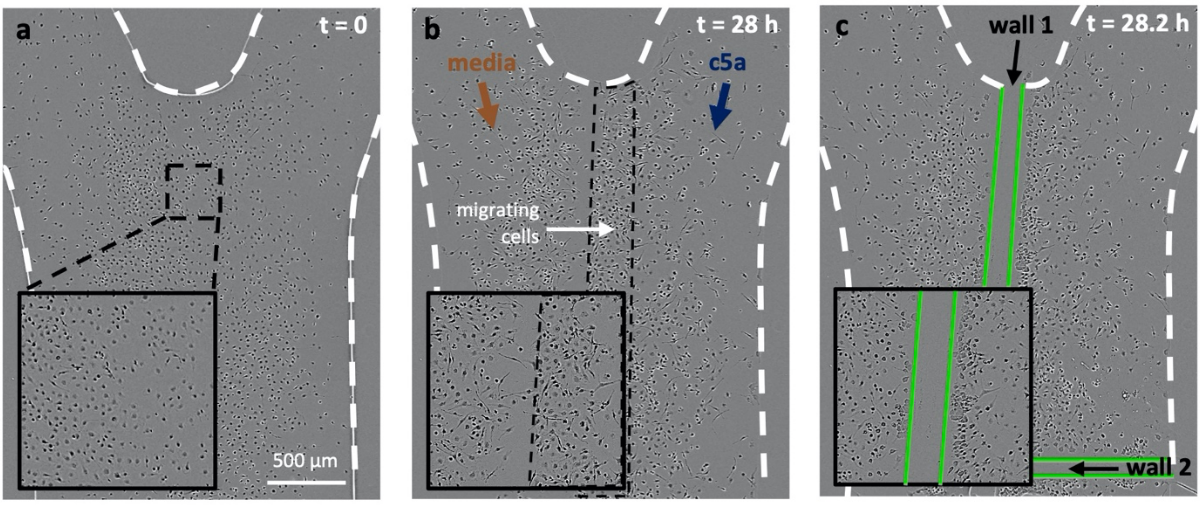


**Figure S8. Reconfiguring a microfluidic circuit on a microscope to isolate populations of chemotaxing BMDMs.** Chemotaxing BMDMs in response to a gradient of C5a are isolated from the population of non-migrating cells using live microscopy, by using an adaptor fitted to the condenser of the microscope (as in **Fig. 2**) **(A)** BMDMs are plated into the central arm of an 'm'-shaped circuit (inset shows close-up of cells). **(B)** Two laminar streams of a chemoattractant (C5a, right) and cell culture media (left) are generated over the cells. Diffusion creates a stable concentration gradient of C5a along the length of the conduit. Cells within the gradient sense the increasing concentration of C5a and migrate towards it over ~28 h. Analysis of the movie made allows us to determine the region containing the population of migrating cells (in the dashed box). **(C)** The microscope is fitted with a similar adaptor to that shown in **Figure 2** which holds a PTFE stylus used to make a new fluid wall along the length of the conduit (green, 'wall 1'). This isolates the population of migrating cells (to the right of wall 1) from non-responsive cells. 'Wall 2' downstream and 'wall 3' upstream (not visible in the image) are then made to completely seal the cells from the rest of the circuit (as in **Fig. 3C**).

### Supplementary Videos

**Supplementary Video 1. Phase contrast movie of chemotaxing macrophages.** BMDMs in a C5a gradient (media input at left, and 10 nM C5a at right); they chemotax towards C5a over ~28 h.

**Supplementary Video 2. Phase contrast movie of chemotaxing bacterial cells.** *P. aeruginosa* in ciprofloxacin gradient (TB + a phase-dark blue dye is input at left to give the dark region, and the antibiotic at 100X MIC is input on the right to give the phase-light region); they chemotax towards the antibiotic over ~24h. The gradient extends over ~90 µm.

### Supplementary STEP files of the 3-D printed parts.

STEP files from the CAD models of the 3-D printed adaptors used to transform a microscope into a reconfiguring tool.
